## Supplementary material for "Fine-grained simulations of the microenvironment of vascularized tumours": Simulation Details

### Supplementary text for “Fine-grained simulations of the microenvironment of vascularized tumours”

#### ABSTRACT

In this little supplementary file to the main manuscript we elaborate on the implementation, hardware and simulation details.

#### 1 Introduction

All source code and parameters to reproduce our simulations are publicly available at GitHub

VBL : https://github.com/thierry3000/VBL
Tumorcode : https://github.com/thierry3000/tumorcode

and distributed with this supplemental material. For the VBL parameters, see the subfolder `parameters` and for Tumorcode parameters, see the subfolder `py/krebsjobs/parameters` within their sources.

The initial vasculature was created with the Tumorcode program[1] `submitVesselgeneration` and the following arguments: `-p default -t 8 -w 1500 -i 1` , see command line help (`-h` option) for their explanation. The resulted initial vasculature comprises a lattice with a constant of 130 $\mu$m and a total size of (1690 $\mu$m x 2080 $\mu$m x 1820 $\mu$m ). For the cancerous modifications, this is subdivided by a factor of 10 such that the tumor vasculature's lattice constant is 13 $\mu$m . Note that the subdivision of a face centered cubic lattice is again a face centered lattice which is an essential feature of our lattice based simulation. As reported in[2,3], the vessel network is superimposed by regular cubic grid taking care of the continuum equations. We chose a constant of 30 $\mu$m .

We chose VBL to operate with a time step of 50 seconds while the minimal time step in Tumorcode is one hour, so the time scales from VBL and Tumorcode are sufficiently different to interleave the two programs. To this end, the VBL software was redesigned with the submodule or library approach in mind that allows single calls execution a complete procedure.

##### 1.1 Finding the nearest vessel

Finding the nearest vessel for every cell is nearly impossible from measured datasets. For our in silico system this is straightforward.

For every vessel, we compute its center of mass and hand it to the ANN library (`https://www.cs.umd.edu/~mount/ANN/`)[4]. The library uses a special storage structure allowing a quick read out of the nearest neighbors for a given query point which is the cell's position in our case.

##### 1.2 Variation of initial seed

In the main document, we report simulations with 3 different tumor seeding positions: one in the center of the simulation domain, one next to an arterial bifurcation and one next to a venous bifurcation. All other parameters stay the same.

For the seeds next to the bifurcation, we report measurements along 3 different lines. For simplification, we label them here as follows:

**case Ap** a parallel line along an arterial bifurcation

**case Vp** a parallel line along a venous bifurcation

**case Vo** an orthogonal line in between an venous bifurcation

| case | sampling line start | sampling line end | seeding point | taken time step |
|---|---|---|---|---|
| Ap | (195,356.5, -424.5) | (195,506.6,-318.4) | (240, 400, -310) | 350,495 |
| Vp | (349,157,-350) | (344,185,-281) | (360, 180, -310) | 350,495 |
| Vo | (350, 248,-327) | (327, 78,-281) | (360, 180, -310) | 350,495 |

**Table 1.** Detailed position of sampled lines.

#### 2 Computing the oxygen content

VBL outputs the amount of oxygen carried per cell in piko gramm (pg). Knowing the cells volume from the radius and the solubility of oxygen, the partial pressure is calculated and used in all our analysis. For the oxygen solubility within the cell we use the value reported in[5].

$$\alpha = 0.003 \frac{\text{ml O2}}{\text{cm}^3\ \text{mmHg}}$$

#### 3 Videos

To highlight the temporal evolution, we visualized the simulation were the spheroid was seeded in the center by means of 2 videos. Each frame is rendered by Tumorcode which provides a direct call to POV-Ray (http://www.povray.org/). The successive images were combined into to movie of 20 frames per second by the ffmpeg tool (http://www.ffmpeg.org/). The movies show the initial vasculature colored by their blood pressure (left color bar) and cut open in a pie like fashion which allows a better view on the developing spheroid in the center. In the file (https://figshare.com/articles/VBL_tumor_PO2/7406180) the cells are color code due to their value of oxygen partial pressure (right colorbar). Note that we can observe the hypoxic rim by pure eye. In the second file (https://figshare.com/articles/VBL_tumor_pH/7406183) , the cells are color coded by their value of pH . The formation of the regions with different values of pH is observed in the presented microenvironment.

#### 4 Hardware

We used nodes of a linux cluster with 28 cores and 128GB memory. The runtime varied from 4 to 6 weeks. This is expected, because of the large number of random events.
